# Computational Gait Retraining from a Single Wearable Sensor: Composite Trajectory Feedback and Multiscale Lower-Limb Adaptation

**DOI:** 10.64898/2026.09.01.748555

**Authors:** Seth Donahue, Patrick Fischer, Zachary Hoegberg, Matthew J. Major

## Abstract

Clinical gait retraining often relies on multi-sensor systems and high-dimensional feedback displays, increasing setup and interpretation demands. We developed a single-IMU visual biofeedback system that computes Lower Limb Trajectory Error (LLTE) in real time from knee-position and shank-orientation trajectories across stance. A synchronized multi-IMU array independently characterized multi-joint responses. Twenty able-bodied adults walked on a treadmill under flexed-knee and extended-knee targets while receiving either corrected or uncorrected LLTE feedback, where the corrected formulation accounted for limb orientation at initial contact. LLTE and stance-phase knee kinematics changed systematically under the flexed-knee target; adaptation toward the extended-knee target was less directionally consistent. The temporal pattern of error reduction did not differ reliably between feedback formulations under either target. Ankle range of motion also changed across training, whereas hip range of motion did not, indicating that the kinematic response was not confined to the targeted knee. In a duration-matched analysis, multiscale multivariate sample entropy (MSMVSE) was higher during feedback-task walking than during normal walking across both targets and feedback formulations; all planned baseline contrasts remained significant after cluster-based permutation testing and Holm correction. No systematic early-to-late change in MSMVSE was detected within feedback trials. A single wearable-derived composite error signal can therefore drive systematic gait modification, while the lower-limb response extends beyond the variable represented in the feedback. MSMVSE distinguished feedback-task walking from normal walking but did not show progressive within-trial convergence, supporting its use as a complementary descriptor of multiscale kinematic structure rather than a direct measure of inter-joint coordination.

## 1. Introduction

Wearable inertial measurement units (IMUs) enable objective measurement of human movement while also supporting real-time biofeedback during rehabilitation, both within and beyond the clinic [1, 2, 3, 4]. However, many gait-feedback systems rely on multi-sensor configurations to estimate joint-level kinematics, increasing equipment requirements, setup time, and calibration burden [5]. A complementary approach is to derive a task-specific feedback variable from minimal instrumentation rather than reconstruct the full lower-limb kinematic state in real time. Building on the framework introduced in [6], we use a single shank-mounted IMU to compute a composite lower-limb trajectory error (LLTE) and provide visual feedback during prescribed flexed- and extended-knee gait modifications. The single sensor therefore serves as the feedback instrument, whereas an independent multi-IMU array is used to quantify the resulting lower-limb kinematics and determine whether adaptation extends beyond the specific variable presented to the participant.

Visual biofeedback is widely used to elicit motor adaptation [7, 8, 9], and wearable sensor systems extend this capability beyond laboratory settings across clinical populations including Parkinson’s disease, cerebral palsy, lower-limb amputation, and knee osteoarthritis [10, 11, 12, 13, 14]. Feedback variables range from discrete temporal measures [13] to composite kinematic scores [15]. Composite variables can reduce the dimensionality of the explicit feedback task by representing several movement features with a single performance signal, while leaving the underlying coordinative solution under-determined [16, 8, 17]. Prior work demonstrates that individuals can modify multiple features of gait simultaneously when guided by a single composite score [7].

To support near-real-time visual biofeedback from a single IMU, we selected *Lower Limb Trajectory Error* (LLTE) as the composite feedback variable. LLTE integrates knee position in the superior–inferior and anterior– posterior directions with sagittal-plane shank angle throughout stance, quantifying deviation from a reference trajectory as a normalized composite trajectory-error magnitude [18]. Originally developed and validated for prosthetic design optimization [18, 19], LLTE has subsequently been estimated from a single leg-mounted IMU during walking [18], supporting its use here as a real-time task-performance signal for visual gait retraining.

LLTE quantifies performance relative to the prescribed gait trajectory, but a scalar task-level error cannot determine whether adaptation is accompanied by broader changes in the temporal organization of lower-limb kinematics. Multiscale multivariate sample entropy (MSMVSE) provides a complementary description by quantifying the regularity of multiple joint-angle trajectories jointly across temporal scales. The framework extends sample entropy [20, 21, 22] through multiscale analysis [23] and multivariate state-space representations [24, 25], allowing changes in the temporal structure of multijoint kinematics to be examined beyond those captured by a scalar task-performance metric [26, 27]. MSMVSE does not identify specific inter-joint coupling relationships in the manner of relative-phase or coupling-angle analyses; rather, it provides a global multiscale description of multijoint kinematic regularity. To the authors’ knowledge, this framework has not previously been applied to motor adaptation during targeted gait retraining, motivating its use here alongside LLTE and conventional kinematic outcomes.

The purpose of the present study was twofold: first, to evaluate LLTE-based visual biofeedback as a task-specific measure for gait retraining; second, to assess broader changes in lower-limb kinematic organization using joint-level kinematics and MSMVSE derived from an independent outcome-assessment IMU array, so that task-specific performance could be distinguished from broader changes in lower-limb movement. We tested the following hypotheses:

1. LLTE values computed during both corrected and uncorrected gait modifications (flexed- and extended-knee walking) will differ significantly from normal baseline walking at 0.80 m s^−1^.
2. LLTE magnitude will change across training blocks, and this adaptation will be moderated by feedback formulation (corrected vs. uncorrected), such that the pattern of change over time differs between groups.
3. Visual biofeedback will produce significant changes in stance-phase knee kinematics, specifically maximum knee flexion, minimum knee flexion and knee range of motion, across training blocks, with group differences expected for the extended-knee but not the flexed-knee target condition.
4. MSMVSE will differ between normal walking and feedback-task walking; an exploratory secondary analysis will test whether the multiscale entropy profile changes from early to late feedback windows.

## 2. Methods

### 2.1. Participants

Twenty healthy young adults participated (10 men, 10 women; age: 25.79 ± 4.53 years; height: 1.71 ± 0.09 m; weight: 73.23 ± 18.37 kg). All participants provided written informed consent prior to enrollment. The study protocol was approved by the Northwestern University Institutional Review Board (STU00220488), and all procedures conformed to relevant ethical guidelines and regulations. Participants reported no neurological or musculoskeletal disorders.

Two participants, one from each feedback group, were identified a priori as having unusable sensor records and were excluded from the LLTE adaptation and MSMVSE analyses. The resulting analysis cohort comprised 18 participants, nine per feedback group. One further participant in the uncorrected group did not complete the fifth flexed-knee exposure; because the mixed models are complete-case, that participant is absent from the flexed-knee LLTE model only, giving 17 complete cases for that target (uncorrected *n* = 8, corrected *n* = 9) against 18 for the extended-knee target (nine per group). The same 18-participant cohort was used for the baseline target comparison (Hypothesis 1). For the kinematic analyses (Hypothesis 3), the number of complete cases varies by outcome because of additional stride-level data rejection, as described below.

### 2.2. Feedback IMU

Real-time visual biofeedback was delivered from a single IMU (XSENS MTw, Movella, Enschede, Netherlands) affixed to the right shank with a hook-and-loop strap, sampled at 100 Hz. Gait events were identified from this sensor using the algorithm described in [28]: initial contact was detected from acceleration and angular velocity thresholds; mid-stance corresponded to the local minimum in anteroposterior acceleration; toe-off was defined as the peak anteroposterior angular velocity; and mid-swing occurred at the mediolateral angular velocity zero-crossing. LLTE was computed in real time from this shank-mounted sensor for every participant, regardless of foot dominance, according to Eq. (1),

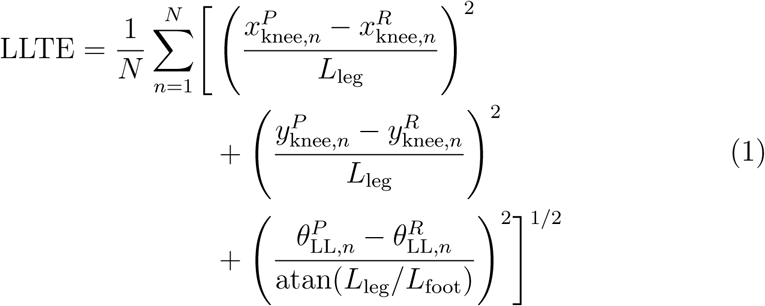

where LLTE is the mean normalized trajectory-error magnitude between measured trajectories (*x^P^*, *y^P^*, *θ^P^*) and reference trajectories (*x^R^*, *y^R^*, *θ^R^*) across the time-normalized stance phase of *N* samples. *x*, *y*, and *θ* denote global superior-inferior position, anterior-posterior position, and shank-to-vertical angle, respectively. Leg length *L*_leg_ was measured from the lateral femoral epicondyle to the floor; *L*_foot_ denotes foot length. At each time-normalized stance sample, the three normalized trajectory errors are combined as a Euclidean magnitude and then averaged across stance; lower LLTE values therefore indicate closer agreement with the prescribed target trajectory. In the real-time implementation, a completed-stance LLTE value was generated stride by stride and the visual display was updated every second initial contact using the mean of the preceding three stance phases. The display also retained the previous five LLTE values as a short visual history. This temporal averaging was used only for feedback delivery; the separate outcome-assessment IMU array did not contribute to the online calculation.

### 2.3. Outcome-Assessment IMU Array

Multi-joint kinematic outcomes (ankle, knee, hip) were characterized using a separate, synchronized eight-sensor IMU array (bilateral foot, shank, and thigh, plus sacrum and xiphoid process), affixed with hook-and-loop straps and sampled at 100 Hz. This array was used exclusively for offline outcome assessment; it did not contribute to the real-time feedback signal described in Section 2.2. Sensor orientation was standardized using static and dynamic calibration procedures [6]. The static calibration consisted of a neutral standing posture held for three seconds. The dynamic calibration comprised several toe-touch trials and treadmill walking at 0.80 m s^−1^, from which offsets between the sensor global coordinate frame and a segment-fixed reference frame were computed; full calibration details are provided in [6].

### 2.4. Experimental Paradigm

Participants completed one minute of treadmill walking at each of 0.80, 1.00, 1.25, and 1.50 m s^−1^ for baseline assessment, followed by two minutes of familiarization with null LLTE biofeedback at 0.80 m s^−1^ (Fig. 1). Participants were instructed to explore knee flexion and extension to minimize displayed error; instructions were standardized as follows: *“This adaptation protocol focuses on the knee joint. You will be asked to adapt to two different targets. Please explore the range of knee flexion and extension to minimize the error between the red bars and the black bar. The red bars of increasing opacity represent the time history of the previous five LLTE values. Adjust your walking pattern gradually, as the feedback is delayed.”* Feedback was updated every second initial contact as the average of the preceding three stance phases, with standardized verbal prompts delivered at 30 s, 1.5 min, and 2.5 min.

**Figure 1:**
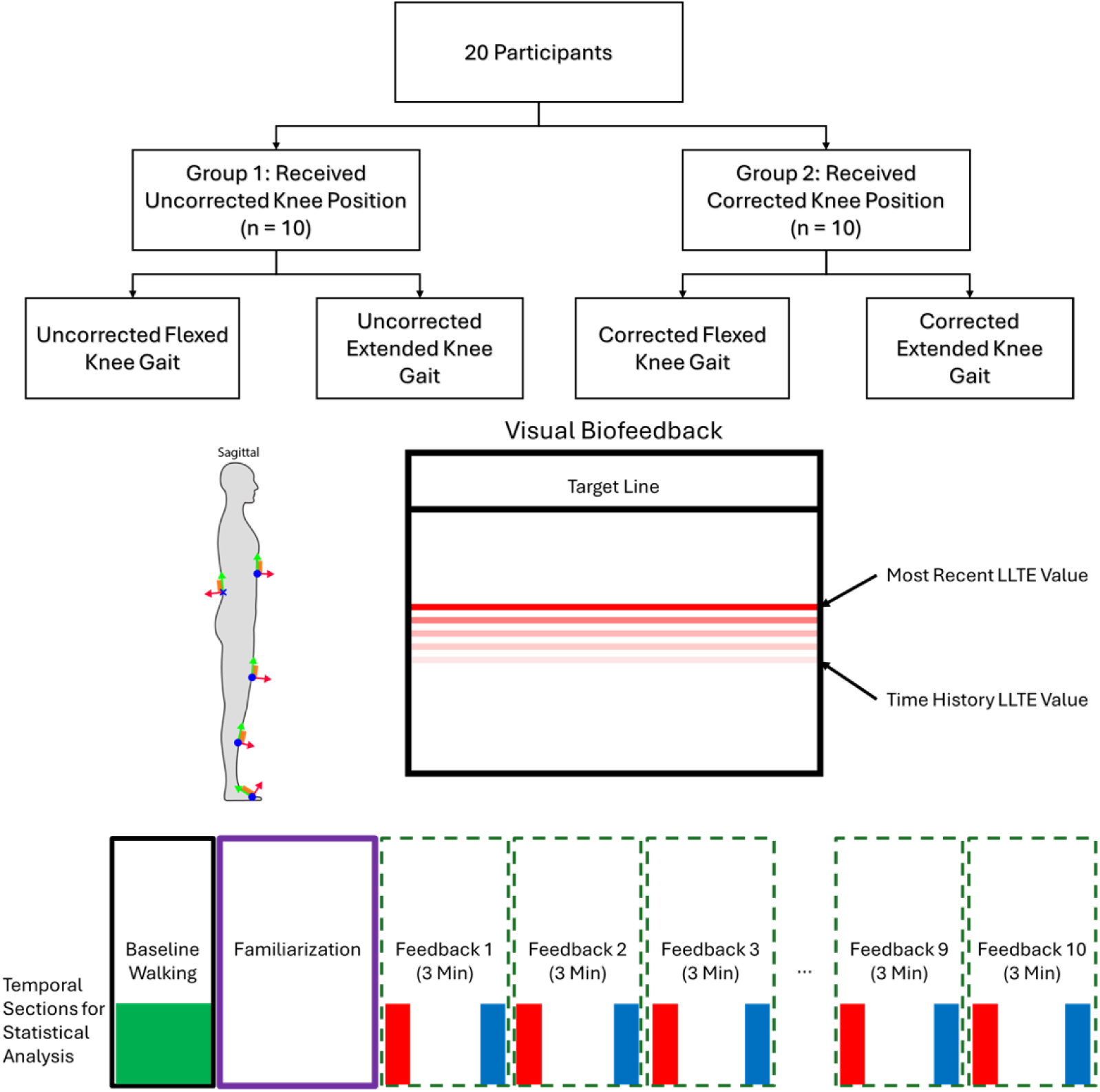
Participants were assigned to one of two groups, each exposed to a distinct reference trajectory and aiming to achieve an LLTE of zero (top). Kinematics were recorded using IMUs placed bilaterally on the feet, shanks, and thighs, with additional sensors on the sacrum and xiphoid process (left). Real-time visual biofeedback displayed the current LLTE value (red), a short temporal history represented by progressively fading bars, and the prescribed target level (black). The experimental paradigm (bottom) comprised baseline walking (normative data collection), familiarization, and ten 3-minute feedback trials, with 5 trials in random order with each target. To quantify learning-related changes, the first and final 30 seconds of each feedback trial were analyzed.

Participants were randomly assigned to one of two feedback groups that differed in how knee trajectory was referenced within the LLTE computation. In the *uncorrected* group, knee trajectory was referenced to a fixed origin regardless of limb orientation at initial contact; consequently, deviations in shank orientation at foot strike contributed to the anterior–posterior knee-position error during stance. In the *corrected* group, the trajectory origin was offset using the sine of shank angle at initial contact, reducing LLTE sensitivity to this orientation-dependent shift. The two formulations therefore presented the same scalar error quantity but differed in whether initial-contact limb orientation was retained within the trajectory mismatch.

Both groups performed two tasks requiring gait modification to match a pre-defined target trajectory reflecting either flexed-knee or extended-knee walking. Each task consisted of five trials presented in randomized order, yielding ten 3-minute feedback trials and introducing contextual interference [29]. Biofeedback was displayed on a monitor positioned in front of the treadmill.

### 2.5. Data Analysis

Real-time IMU data were processed using the ReBAIT open-source framework [6]. Sagittal-plane ankle, knee, and hip angles were derived from the outcome-assessment IMU array, gap-filled by cubic interpolation, and filtered using a bidirectional fourth-order low-pass Butterworth filter with a 24 Hz cutoff [30]. Stance phases longer than 1 s and strides containing joint angles outside physiologic bounds (*> |*180°|), attributable to quaternion-to-Euler conversion artifacts or calibration failure, were excluded from the relevant kinematic analyses.

A normative LLTE reference was established from ensemble-averaged knee anterior–posterior displacement, knee superior–inferior displacement, and shank-angle trajectories during treadmill walking at 0.80 m s^−1^, following [31]. To preserve fidelity with the real-time implementation, the target trajectories used by the feedback calculation were not subsequently filtered or gap-filled. LLTE was evaluated relative to this reference during baseline walking and across feedback trials. Motor adaptation was characterized from the first and final 30 s of each training block using LLTE and stance-phase joint kinematics, with maximum knee flexion specified as the primary kinematic outcome.

Discrete joint-angle outcomes were extracted over the middle 80% of stance, excluding the initial and terminal 10% of the stance phase to avoid the heel-strike and toe-off transitions at which quaternion-to-Euler artifacts are most pronounced. For each stride within an analysis window, the maximum, minimum and range of the joint angle were taken over that interval, and the resulting per-stride values were averaged across all valid strides in the window to yield one observation per participant, joint, target and block. These discrete kinematic outcomes were computed for the right limb, which carried the feedback sensor; the multiscale entropy analysis described below instead used all six bilateral joint-angle trajectories.

### 2.6. Computational Analysis of Multiscale Kinematic Structure

Multiscale multivariate sample entropy (MSMVSE) was calculated from six sagittal-plane joint-angle trajectories: bilateral hip, knee, and ankle angles. MSMVSE was used to characterize the regularity of multijoint kinematics across temporal scales and was not interpreted as a direct measure of inter-joint coordination. For each participant, normal walking at 0.80 m s^−1^ was compared with the first and final feedback exposures for each target condition. To avoid confounding entropy estimates by unequal record length, feedback windows were duration-matched to each participant’s available normal-walking baseline (57.35–60.00 s). The earliest continuous segment of each feedback trial was extracted for the early-window analysis, and an equivalently sized segment ending at the final recorded sample was extracted for the late-window analysis. All six trajectories were interpolated to 100 Hz before entropy calculation.

Sample entropy used embedding dimension *m* = 2 and a tolerance of 0.20 times the pooled standard deviation of the six signals within each analysis window. The resulting tolerance was held fixed across scales within that window. Scales *τ* = 1–16 constituted the common finite range for comparisons with normal walking. Early-versus-late feedback comparisons were restricted to scales *τ* = 1–15 because one late-window estimate was undefined at scale 16. The original full-duration, 20-scale MSMVSE curves were retained as a sensitivity analysis.

### 2.7. Statistical Analysis

All analyses used a critical alpha of 0.05. Normality was assessed using the Kolmogorov–Smirnov test. LLTE was right-skewed and was log-transformed prior to analysis; the discrete joint-angle outcomes did not require transformation and were analysed on their original scale, in degrees.

#### Hypothesis 1

A one-way repeated-measures ANOVA (*n* = 18) compared LLTE values at baseline across five conditions: normal walking at 0.80 m s^−1^ and the four target formulations (corrected and uncorrected flexed- and extended-knee), each evaluated using the 0.80 m s^−1^ walking data. Post hoc comparisons used Bonferroni-corrected paired *t*-tests across the full set of ten pairwise condition contrasts; the four contrasts against normal walking are reported below. Because the correction was applied across all ten contrasts rather than only the four against the control condition, these post hoc tests are conservative relative to a control-referenced procedure such as Dunnett’s test.

#### Hypothesis 2

Adaptation effects were evaluated using separate two-way mixed ANOVAs for the flexed- and extended-knee targets. Each model included a within-subject factor of *Block* (eleven levels: baseline plus the first and final 30 s of each of five 3-min training blocks) and a between-subject factor of *Group* (Corrected vs. Uncorrected), with log-transformed LLTE magnitude as the dependent variable. The extended-knee model comprised 18 complete cases (nine per group) and the flexed-knee model 17 complete cases (uncorrected *n* = 8, corrected *n* = 9), as described above. Significant main effects or interactions were followed by Bonferroni-adjusted pairwise comparisons. Sphericity of the within-subject factor was assessed with Mauchly’s test. It was violated for the extended-knee model (*W* = 0.0003, *χ*^2^(54) = 109.9, *p <* 0.001) but not for the flexed-knee model (*W* = 0.005, *χ*^2^(54) = 66.2, *p* = 0.171). Greenhouse–Geisser corrected degrees of freedom are therefore reported for all within-subject effects of the extended-knee model, and are additionally reported for the flexed-knee model for comparability. The correction factor was applied to the degrees of freedom of the mixed model itself, as df_1_ = (*k −* 1)*ε* and df_2_ = (*k −* 1)(*N − g*)*ε*, and the same *ε* was applied to the Group *×* Block interaction.

#### Hypothesis 3

Maximum knee flexion, minimum knee flexion, knee range of motion (ROM), ankle ROM, and hip ROM were examined using the same Block *×* Group mixed-ANOVA structure, separately for each target condition, using the stride-averaged middle-80%-of-stance outcomes defined above. Sphericity was assessed per model with Mauchly’s test and Greenhouse– Geisser corrections were applied where it was violated. These models are complete-case over the eleven Block levels, and the number of complete cases varies across outcomes because of stride-level data rejection, from 16 to 18 participants. Neither participant with a prespecified bad-data exclusion contributes a complete series to any kinematic outcome, so all ten models are unchanged whether or not those exclusions are applied, and the kinematic cohort is consistent with that used for the LLTE and MSMVSE analyses.

#### Hypothesis 4

Paired differences in MSMVSE were tested across adjacent temporal scales using two-sided cluster permutation tests. All participant-level sign flips were enumerated. Adjacent scales showing effects in the same direction were grouped into clusters, cluster mass served as the test statistic, and the largest cluster mass observed in each permutation formed the family-wise null distribution across scales. Holm correction was then applied across the eight planned contrasts within each analysis family. The primary baseline analysis compared duration-matched normal-walking and feedback-task windows. A secondary early-versus-late analysis tested for within-trial change in the MSMVSE profile. For descriptive effect estimation, mean paired differences and 95% confidence intervals were also reported. The original full-duration, 20-scale analysis was retained only as a sensitivity analysis.

## 3. Results

### 3.1. LLTE Target Comparison During Normal Walking

To examine how LLTE values associated with each biofeedback target differed from normal gait, LLTE was compared across five conditions during treadmill walking at 0.80 m s^−1^: four visual biofeedback targets (corrected and uncorrected flexed- and extended-knee) and a normal walking control. The control condition represented the aggregate LLTE trajectory computed across the 18 analysed participants during normal walking at 0.80 m s^−1^. LLTE values for each target condition were extracted from the corresponding 0.80 m s^−1^ walking bouts (Fig. 2).

**Figure 2:**
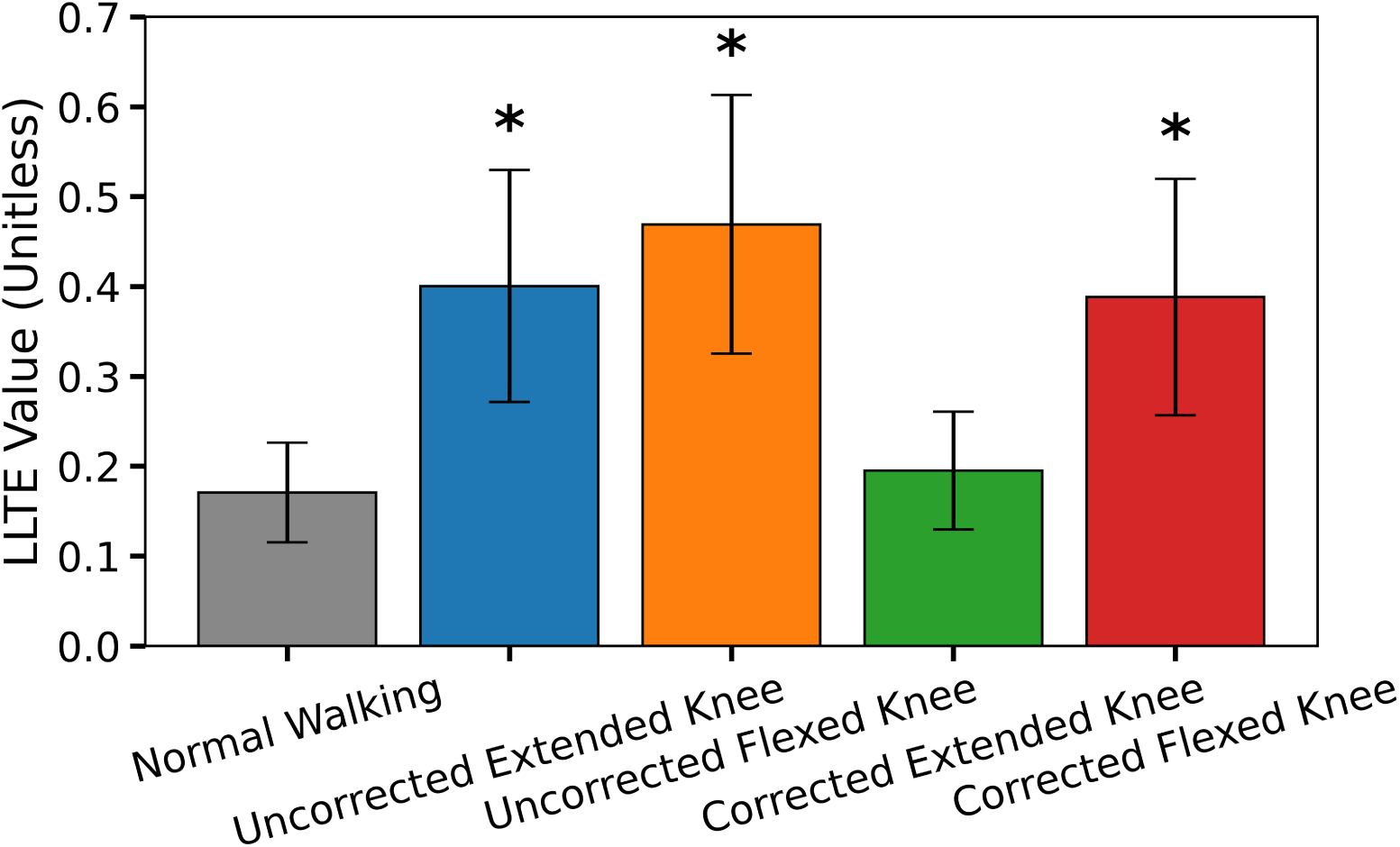
Mean LLTE values during normal walking (0.80 m s^−1^) compared across four biofeedback target conditions. The gray bar represents the aggregate normal walking mean LLTE value, i.e., the expected baseline LLTE. Blue and orange bars indicate uncorrected feedback targets (extended- and flexed-knee, respectively), and green and red bars indicate corrected feedback targets (extended- and flexed-knee, respectively). Error bars denote ±1 SD. Asterisks indicate significant differences from the normal walking condition (Bonferroni-corrected paired *t*-tests across all ten pairwise contrasts, *p <* 0.001); the corrected extended-knee target did not differ significantly from normal walking.

There was a significant main effect of target condition on LLTE values. Mauchly’s test returned numerically invalid output (sphericity statistic and *χ*^2^ outside their valid range), consistent with a near-singular covariance matrix arising from near-collinearity between the two flexed-knee target conditions (*r* = 0.996); sphericity was therefore treated as violated and a Greenhouse–Geisser correction was applied manually (*ε* = 0.288, near the theoretical floor of 1*/*(*k −* 1) = 0.25), *F* (1.15, 19.59) = 103.74, *p* = 1.2 *×* 10^−9^, 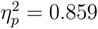 (Fig. 2). Bonferroni-corrected post hoc comparisons showed that LLTE values differed significantly from normal walking for the uncorrected flexed-knee (*p <* 0.001), uncorrected extended-knee (*p <* 0.001), and corrected flexed-knee targets (*p <* 0.001), whereas LLTE values for the corrected extended-knee target did not differ significantly from normal walking (*p* = 0.093).

This last comparison should be interpreted cautiously. Its uncorrected value was *p* = 0.009, so the non-significant result reflects correction across all ten pairwise contrasts; a control-referenced procedure restricted to the four contrasts against normal walking would place it below the 0.05 threshold (*p* = 0.037). The conclusion that the corrected extended-knee target lies closest to habitual gait therefore rests on the ordering of effects rather than on a categorical significance boundary.

### 3.2. LLTE Biofeedback

LLTE values for the extended-knee condition, including baseline and the first and last 30 seconds of each adaptation trial, are shown in Fig. 3. The main effect of Group was not significant, *F* (1, 16) = 2.70, *p* = 0.120, 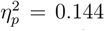. The main effect of Block was significant (Greenhouse–Geisser corrected, *ε* = 0.49), *F* (4.89, 78.28) = 2.98, *p* = 0.017, 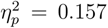. The Group *×* Block interaction was not significant once the same correction was applied, *F* (4.89, 78.28) = 2.08, *p* = 0.079, 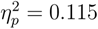; it reached nominal significance only in the uncorrected test (*p* = 0.029), and sphericity was clearly violated in this model, so it is not interpreted as a reliable group difference.

**Figure 3:**
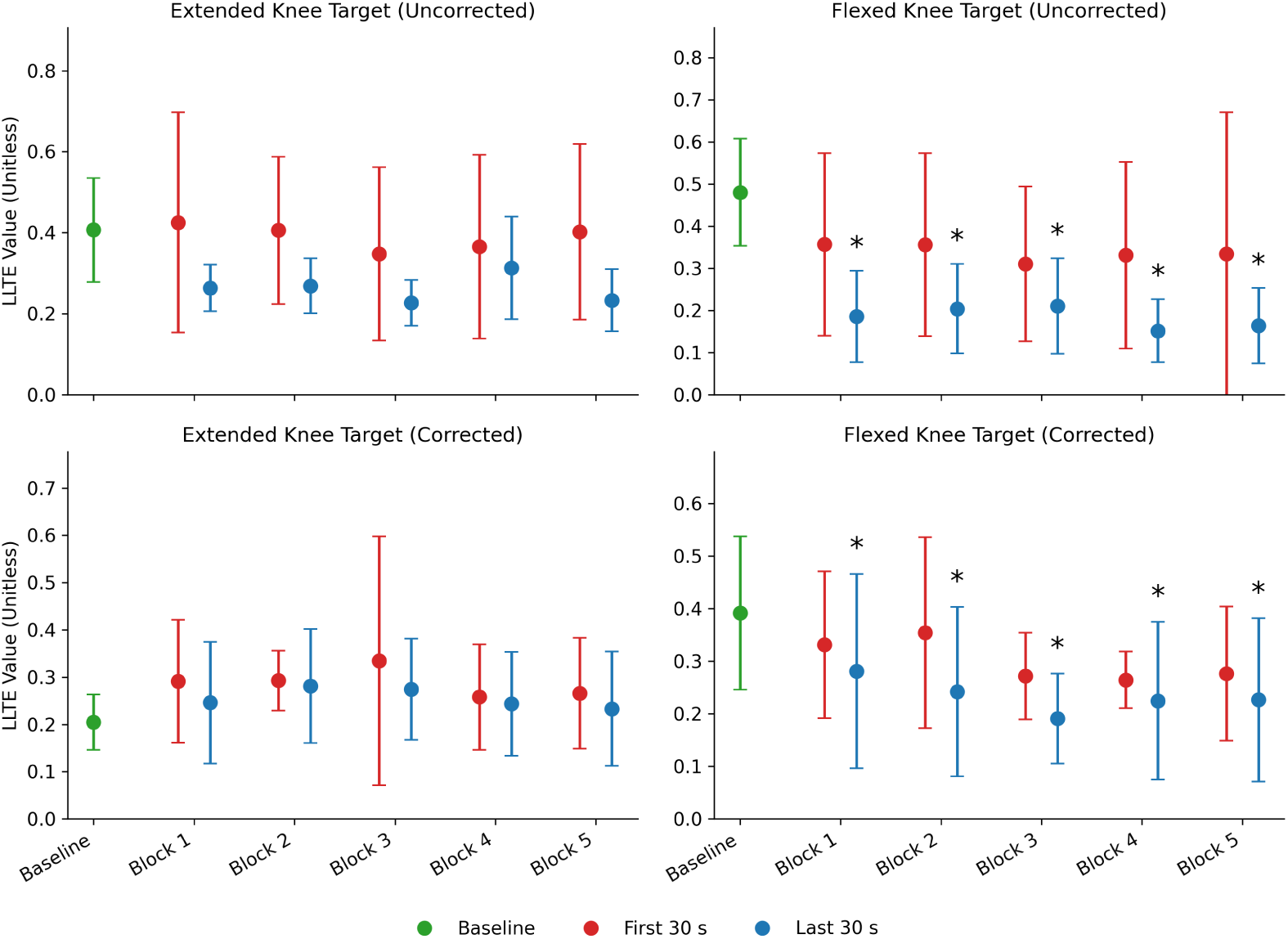
Aggregate LLTE (mean ± SD) during baseline and the first and last 30 seconds of visual-biofeedback-driven adaptation, across training blocks for extended-knee (left panels) and flexed-knee (right panels) target conditions. Smaller LLTE values indicate closer agreement with the target trajectory. Green markers represent baseline LLTE during normal walking at 0.80 m s^−1^ prior to biofeedback training; red and blue markers represent the first and last 30 s of each training block, respectively. Uncorrected feedback groups are shown in the top panels and corrected feedback groups in the bottom panels. Asterisks denote Bonferroni-corrected contrasts between baseline and the last 30 s of a training block that were significant in the Block main effect, which is pooled across feedback groups because the Group *×* Block interaction was not significant for either target; the same markers therefore apply to both panels of a column. No baseline contrast reached significance for the extended-knee target.

A follow-up comparison of the two feedback groups within each block level clarifies the origin of the uncorrected interaction. The groups differed significantly only at baseline, *t*(16) = 4.46, *p* = 0.004 (Bonferroni-corrected), *η*^2^ = 0.525, with no significant group difference at any of the ten subsequent training-block levels (all corrected *p ≥* 0.92). The uncorrected interaction therefore reflects pre-existing baseline non-equivalence that converged once feedback began, rather than differential adaptation to the two feedback formulations. Among block-level pairwise comparisons, only the contrast between the first 30 s of Block 2 and the last 30 s of Block 5 was significant (*p* = 0.026, *η*^2^ = 0.238); no comparison against baseline survived correction for this target.

For flexed-knee gait, Mauchly’s test did not indicate a departure from sphericity. The main effect of Group was not significant, *F* (1, 15) = 0.16, *p* = 0.698, 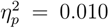 and neither was the Group *×* Block interaction, *F* (10, 150) = 1.49, *p* = 0.148, 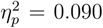 (*ε* = 0.55; *F* (5.52, 82.73) = 1.49, *p* = 0.197 when the same correction is applied). The main effect of Block was significant, *F* (10, 150) = 14.07, *p <* 0.001, 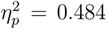, and remained so under Greenhouse–Geisser correction, *F* (5.52, 82.73) = 14.07, *p <* 0.001. Error reduction under the flexed-knee target was therefore systematic across the training session and did not depend on feedback formulation.

Bonferroni-corrected paired comparisons of each block’s own first-versus-last 30 s segment were significant for Block 1 (*p* = 0.048), Block 2 (*p* = 0.010), and Block 3 (*p* = 0.030), but not for Block 4 (*p* = 0.080) or Block 5 (*p* = 0.104); within-block LLTE reduction was therefore confirmed in three of five training blocks, concentrated in the earlier part of the session. A broader pattern of significant reduction was evident in cross-block comparisons against baseline and against earlier blocks’ first-30 s segments, summarized in Table 2. LLTE at the last 30 s of every training block was significantly lower than baseline. One contrast ran in the opposite direction: LLTE rose significantly between the last 30 s of Block 3 and the first 30 s of Block 4 (*p* = 0.050, *η*^2^ = 0.164), consistent with partial between-block washout under the randomized, contextually interfering trial order.

### 3.3. Lower Extremity Kinematic Patterns

Kinematic analyses of the knee joint (Fig. 4) showed significant within-subject Block effects for maximum and minimum stance-phase knee flexion angle in both target conditions. For knee flexion maxima, Block effects were significant for both the extended-knee target (Greenhouse–Geisser corrected, *ε* = 0.44, *F* (4.39, 65.84) = 4.12, *p* = 0.005, 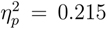), and the flexed-knee target (*ε* = 0.47, *F* (4.73, 66.27) = 19.41, *p <* 0.001, 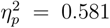).

**Figure 4:**
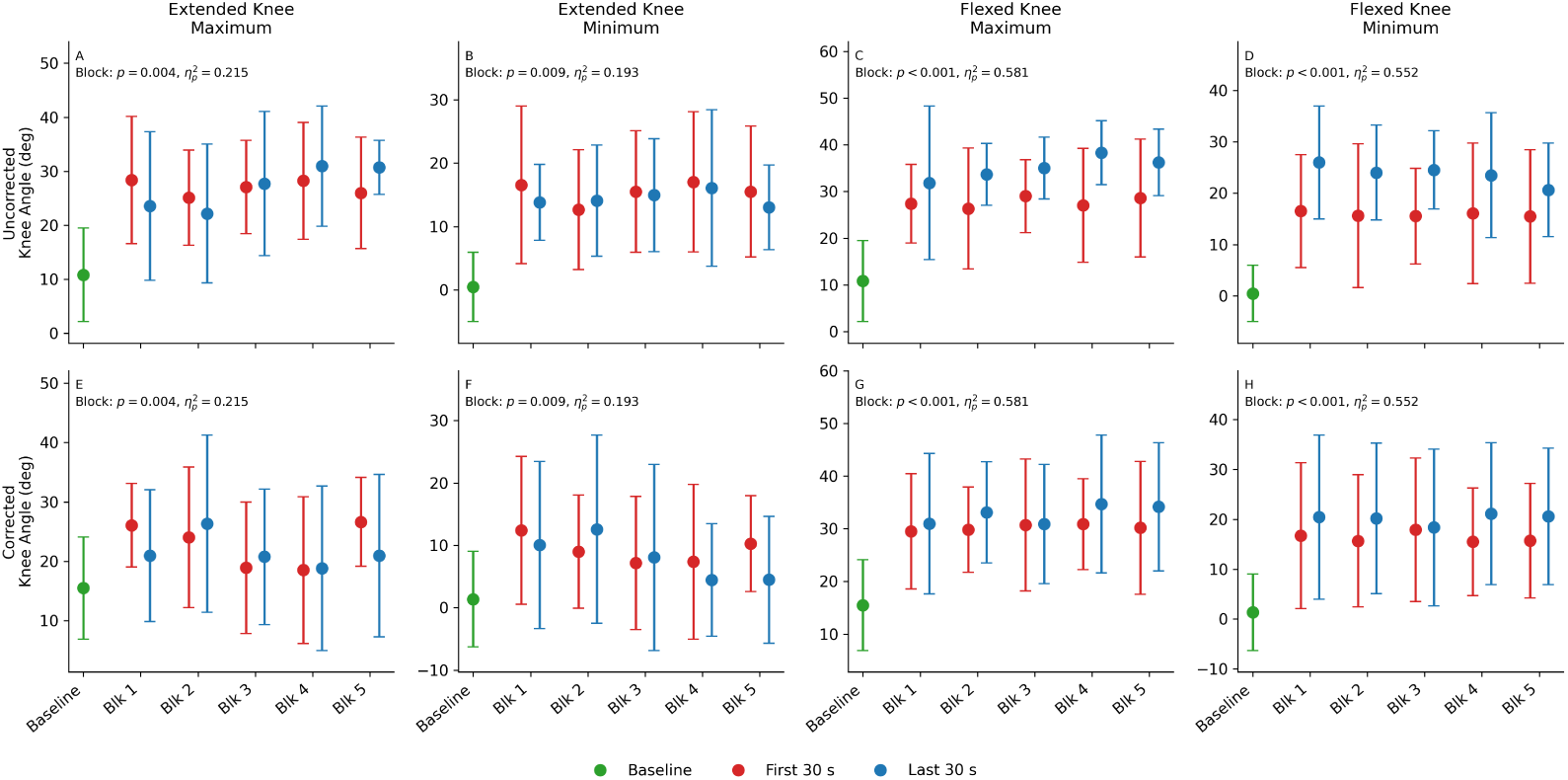
Stance-phase knee kinematics (mean ± SD) during baseline walking and across the five biofeedback training blocks, taken over the middle 80% of stance on the right limb. Columns differentiate target conditions (extended-knee vs. flexed-knee gait) and discrete gait parameters (maximum vs. minimum knee flexion angle). Rows distinguish the uncorrected (panels A–D) and corrected (panels E–H) feedback groups. Green markers represent unadapted baseline kinematics during normal walking at 0.80 m s^−1^. Red and blue markers denote the first and last 30 s of each training block, respectively. Each panel is annotated with the main effect of Block for that outcome and target, Greenhouse–Geisser corrected where sphericity was violated; no post hoc pairwise comparisons were performed for the knee kinematic outcomes.

Minimum knee flexion showed a similar pattern: extended-knee (*ε* = 0.43, *F* (4.28, 64.24) = 3.58, *p* = 0.010, 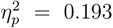); flexed-knee (*ε* = 0.48, *F* (4.81, 67.39) = 17.23, *p <* 0.001, 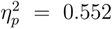). Knee ROM also varied significantly by Block for both targets. Sphericity was not violated for the extended-knee target, so uncorrected degrees of freedom are reported, *F* (10, 150) = 15.29, *p <* 0.001, 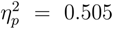; for the flexed-knee target a Greenhouse–Geisser correction was applied (*ε* = 0.52), *F* (5.20, 78.06) = 14.46, *p <* 0.001, 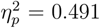. A between-subject Group effect was observed only for knee ROM in the extended-knee target condition, *F* (1, 15) = 6.36, *p* = 0.023, 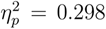; the corresponding flexed-knee effect did not reach significance, *F* (1, 15) = 4.01, *p* = 0.064, 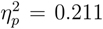. All remaining Group effects were non-significant (all *p ≥* 0.082), as was every Block *×* Group interaction (all *p ≥* 0.075 uncorrected, and *p ≥* 0.144 after Greenhouse–Geisser correction).

For ankle ROM (Table 1), Block effects were significant for both targets: extended-knee (*ε* = 0.47, *F* (4.75, 75.94) = 11.87, *p <* 0.001, 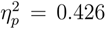); flexed-knee (*F* (10, 150) = 12.62, *p <* 0.001, 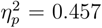), with no significant Group or interaction effects (all *p ≥* 0.54). Hip ROM (Table 1) did not vary significantly across Blocks: extended-knee (*ε* = 0.51, *F* (5.14, 77.10) = 1.10, *p* = 0.381, 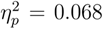); flexed-knee (*ε* = 0.42, *F* (4.20, 62.94) = 1.74, *p* = 0.141, 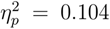); no Group or interaction effects were significant (*p ≥* 0.187).

**Table 1:** Mean (±SE) Ankle and Hip Range of Motion (Degrees, Middle 80% of Stance) at Baseline and During Early (Block 1) and Late (Block 5) Biofeedback Adaptation (Last 30 s). Values are group means with standard errors. These descriptives are computed over all participants contributing valid strides to each cell rather than over the complete-case cohort of the corresponding ANOVA, so the number of contributing participants varies by cell.

|  | **Ankle ROM** |  | **Hip ROM** |  |
|---|---|---|---|---|
| **Block** | **Uncorr.** | **Corr.** | **Uncorr.** | **Corr.** |
| *Flexed-Knee Target* |  |  |  |  |
| Baseline | 11.53 ± 1.84 | 14.75 ± 3.09 | 35.81 ± 2.17 | 38.33 ± 2.44 |
| Early (Blk 1) | 21.07 ± 2.17 | 21.33 ± 2.05 | 40.51 ± 2.74 | 34.25 ± 3.57 |
| Late (Blk 5) | 20.17 ± 1.88 | 22.07 ± 1.49 | 40.81 ± 2.46 | 41.62 ± 4.07 |
| *Extended-Knee Target* |  |  |  |  |
| Baseline | 11.53 ± 1.84 | 14.75 ± 3.09 | 35.81 ± 2.17 | 38.33 ± 2.44 |
| Early (Blk 1) | 22.40 ± 1.84 | 24.09 ± 2.11 | 36.89 ± 4.00 | 34.16 ± 2.97 |
| Late (Blk 5) | 20.98 ± 2.32 | 26.21 ± 2.07 | 40.86 ± 2.60 | 39.08 ± 4.73 |

### 3.4. Multiscale Multivariate Sample Entropy

Eighteen participants contributed to the duration-matched MSMVSE analysis, with nine participants in each feedback group. Across both feedback formulations and both gait targets, MSMVSE was higher during feedback-task walking than during normal walking at 0.80 m s^−1^ (Fig. 5). This pattern was present in both early and late feedback windows. All eight planned feedback-versus-baseline contrasts remained significant after cluster-based inference across scales and Holm correction (*p* = 0.031–0.047). Averaged descriptively across the examined scales, feedback-minus-baseline differences ranged from approximately 0.21 to 0.36 MSMVSE units. For reference, normal-walking MSMVSE means ranged from 0.09 at scale 1 to 0.46 at scale 16, so these differences are comparable in magnitude to the baseline values themselves rather than representing a small perturbation of them. The original full-duration, 20-scale sensitivity analysis showed the same broad pattern.

**Figure 5:**
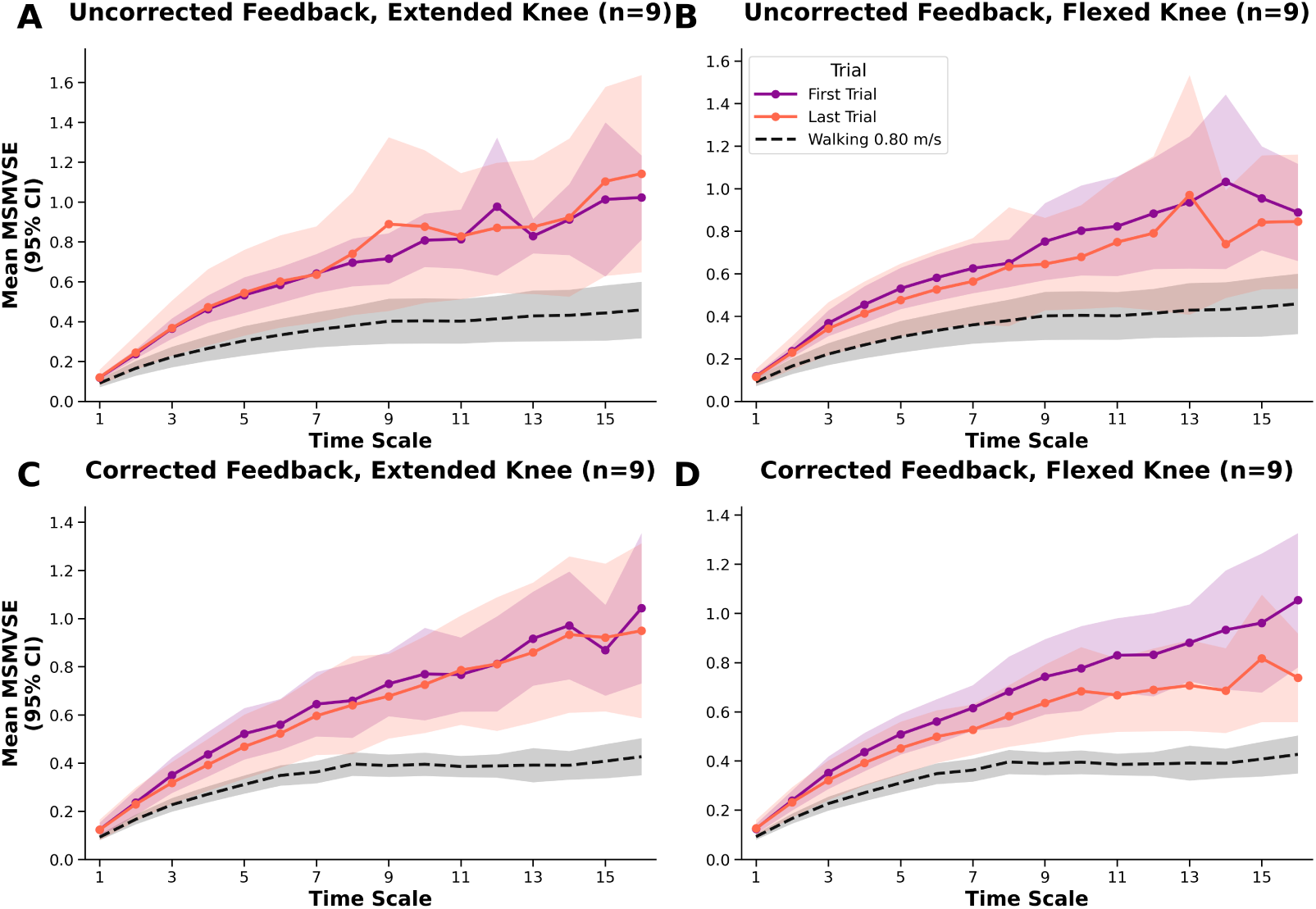
Duration-matched MSMVSE profiles for normal walking and feedback-task walking across corrected and uncorrected flexed- and extended-knee target conditions. Lines show mean MSMVSE across temporal scales, with uncertainty bands illustrating within-condition variability. Inferential testing was performed on paired differences using cluster permutation tests across adjacent scales, followed by Holm correction across the eight planned contrasts within each analysis family.

No systematic early-to-late change in the MSMVSE profile was detected within feedback trials. No significant early-versus-late cluster formed in any group-by-target condition. Mean late-minus-early differences ranged from *−*0.036 to 0.075 MSMVSE units, and all corresponding 95% confidence intervals included zero. Thus, the multiscale entropy difference relative to normal walking was already evident early in the feedback task, whereas the present analysis did not detect progressive within-trial change.

Late feedback windows also remained elevated relative to normal walking in the primary analysis. Because two final trials were shorter than the duration required to obtain completely independent early and late windows, their early and late windows overlapped by 18–19 s; a sensitivity analysis therefore excluded those overlapping window pairs. Six of the eight late-window baseline contrasts remained significant after this exclusion. The two corrected-feedback final-exposure contrasts no longer survived Holm correction (*p* = 0.078), indicating that evidence for persistent late-window elevation under corrected feedback was less robust to window independence than the corresponding findings in the other conditions.

## 4. Discussion

A single composite kinematic variable computed from one shank-mounted IMU was sufficient to systematically modify gait pattern in able-bodied adults, with the clearest response observed for the flexed-knee target. The real-time signal compressed knee-position and shank-orientation trajectories into a scalar task error, while a separate multi-IMU array showed that the resulting kinematic response was not confined to the knee. Knee and ankle ROM changed across training, whereas hip ROM did not. These findings support the use of low-dimensional wearable feedback to drive targeted gait modification while also illustrating that a scalar feedback variable does not fully describe the resulting lower-limb response.

**Table 2:** Significant Bonferroni-corrected pairwise comparisons for LLTE during flexed-knee gait (log-transformed), pooled across both feedback groups (*N* = 17). Cells show *p* (Bonf.) / *η*^2^. Only significant comparisons are shown. The Group *×* Block interaction was not significant for this target, so a single pooled set of block contrasts is reported.

| **vs. Last 30 s of:** | **Blk 1** | **Blk 2** | **Blk 3** | **Blk 4** | **Blk 5** |
|---|---|---|---|---|---|
| Baseline | 0.024 / 0.349 | 0.005 / 0.400 | <0.001 / 0.460 | <0.001 / 0.494 | <0.001 / 0.465 |
| Blk 1 — first 30 s | 0.048 / 0.152 | 0.003 / 0.183 | 0.012 / 0.248 | 0.005 / 0.292 | 0.006 / 0.267 |
| Blk 2 — first 30 s | — | 0.010 / 0.154 | 0.019 / 0.212 | 0.003 / 0.253 | 0.016 / 0.232 |
| Blk 3 — first 30 s | — | — | 0.030 / 0.141 | — | 0.049 / 0.164 |
| Blk 4 — first 30 s | — | — | — | — | 0.029 / 0.187 |

Participants showed the most consistent LLTE and knee-kinematic changes under the flexed-knee target, where error reduction across blocks was large and survived sphericity correction. Contrary to Hypothesis 2, however, the temporal pattern of error reduction did not differ reliably between feedback formulations under either target. The flexed-knee interaction was not significant, and although the extended-knee interaction reached nominal significance in the uncorrected test (*p* = 0.029), sphericity was clearly violated in that model and the effect fell away under Greenhouse–Geisser correction (*p* = 0.079). Follow-up comparisons localized the apparent extended-knee interaction to a single significant group difference at baseline, with no group difference at any training block, indicating convergence from unequal starting points rather than differential learning. Accounting for initial-contact limb orientation within the LLTE computation therefore did not measurably change either the magnitude or the trajectory of adaptation in this sample. Given the modest group sizes, this should be read as an absence of detectable moderation rather than as evidence that the two formulations are equivalent.

The extended-knee target produced a different response. The corrected extended-knee target was already close to participants’ habitual gait pattern at baseline, and subsequent changes were not consistently directed toward the intended extended-knee stance pattern; no block contrast against baseline survived correction for this target. Two features of the task may explain this result. First, LLTE is a scalar magnitude and therefore indicates how far the current trajectory lies from the target without specifying the direction of correction [32]. Second, healthy adults have limited additional knee extension available near terminal stance, reducing the movement space available for further correction. Together, these constraints make the extended-knee target more difficult to achieve using a non-directional scalar error signal.

Within the flexed-knee target, within-block error reduction was concentrated in the first three training blocks, while cross-block comparisons showed that performance at the end of every block remained below baseline. One contrast indicated a significant rise in error between the end of Block 3 and the start of Block 4, consistent with the partial between-block washout expected under a randomized, contextually interfering trial order. Together these patterns suggest rapid early within-block gains that consolidate into a sustained offset from baseline, rather than continuous improvement across the whole session.

Changes outside the knee further clarify what the LLTE signal did and did not control. Ankle ROM changed significantly across both target conditions despite the absence of ankle-specific feedback, whereas hip ROM did not. This pattern demonstrates a kinematic response beyond the knee but does not establish a specific coordination strategy: joint-by-joint ROM cannot determine whether ankle changes were mechanically coupled to knee adaptation or occurred independently. Direct coupling measures such as continuous relative phase or vector coding would be required to make that distinction.

The duration-matched MSMVSE analysis provided a complementary view of the task response. Across both targets and feedback formulations, feedback-task walking showed higher multiscale entropy than normal walking, and all planned baseline contrasts survived cluster-based inference across scales and Holm correction. Matching the analysis windows removes unequal record length as an explanation for this difference. In contrast, no significant early-to-late cluster was detected within feedback trials. The strongest interpretation is therefore that the multiscale entropy profile distinguished the altered feedback-task state from normal walking, not that MSMVSE tracked progressive within-trial adaptation or convergence. A nonsignificant early-to-late result should not be interpreted as equivalence between early and late behavior.

This distinction also limits causal interpretation. Baseline and feedback trials differed not only in the presence of visual biofeedback but also in the prescribed movement target and associated task demands. Higher MSMVSE may therefore reflect target-directed exploration, altered gait mechanics, augmented feedback, or a combination of these factors. MSMVSE describes multivariate signal regularity across scales; it does not identify the joint-to-joint coupling that generated the profile. As reported above, the late-window result was also somewhat sensitive to window independence under corrected feedback, although the broad task-versus-baseline difference was consistent with the original full-duration sensitivity analysis.

Several limitations warrant consideration. First, this study evaluated single-session treadmill adaptation; retention and transfer to overground walking remain untested. Second, the absence of a no-feedback target-matched control condition prevents isolation of visual biofeedback from the effects of target-directed gait modification itself. Third, LLTE is non-directional, which likely limited correction toward the extended-knee target. Fourth, calibration error and quaternion-to-Euler conversion artifacts reduced usable data for some kinematic analyses, and the number of complete cases varies across kinematic outcomes. Fifth, the analysis cohort of 18 participants, reduced to 17 complete cases for the flexed-knee LLTE model, provides limited power to detect moderation by feedback formulation; the absence of a reliable Group *×* Block interaction should be read accordingly. Sixth, groups were not equivalent at baseline under the extended-knee target, which complicates interpretation of between-group comparisons for that condition. Seventh, the MSMVSE analysis was based on a modest sample, coarse scales with undefined estimates were excluded from the matched analysis, and two final trials produced 18–19 s of overlap between early and late windows. Finally, neither MSMVSE nor joint-specific ROM directly quantifies inter-joint coordination. Future work should combine the present real-time scalar feedback framework with direct coupling measures and should test clinical populations in whom gait retraining is an intended therapeutic application.

## 5. Conclusion

A composite trajectory error computed from a single wearable IMU was sufficient to drive systematic gait modification in able-bodied adults, with the most consistent task-specific adaptation observed under the flexed-knee target. Accounting for initial-contact limb orientation within the error computation did not measurably alter the magnitude or the temporal pattern of adaptation in this sample. Independent multi-IMU measurements showed concurrent changes in knee and ankle kinematics, indicating that the lower-limb response extended beyond the variable explicitly represented in the feedback. Duration-matched MSMVSE distinguished feedback-task walking from normal walking across both targets and feedback formulations, but did not show detectable early-to-late change within feedback trials. These findings support LLTE as a low-dimensional real-time feedback variable while defining an important boundary on its interpretation: a scalar task error can guide gait modification without fully characterizing the multijoint dynamics used to achieve it. Multiscale entropy is therefore most informative here as a complementary description of task-related kinematic structure rather than as a direct measure of coordination or learning progression.

## CRediT authorship contribution statement

**Seth Donahue:** Conceptualization, Methodology, Software, Formal analysis, Investigation, Data curation, Visualization, Writing – original draft, Writing – review & editing, Project administration, Funding acquisition.

**Patrick Fischer:** *[Confirm contribution roles]*.

**Zachary Hoegberg:** Methodology, Software, Validation, Writing – review & editing.

**Matthew J. Major:** Conceptualization, Methodology, Supervision, Writing – review & editing.

## Data availability

The data supporting the findings of this study are available through the openICPSR repository at openICPSR Project 212941. The real-time feedback software is built on the open-source ReBAIT framework [6].

## Declaration of generative AI and AI-assisted technologies in the writing process

During the preparation of this work the authors used Claude (Anthropic) in order to edit manuscript text for clarity. The authors reviewed and edited the content as needed and take full responsibility for the content of the published article.

